# Pirfenidone attenuates lung fibrotic fibroblast-mediated fibrotic responses to transforming growth factor-β1

**DOI:** 10.1101/400978

**Authors:** Jin Jin, Shinsaku Togo, Kotaro Kadoya, Miniwan Tulafu, Yukiko Namba, Moe Iwai, Junko Watanabe, Kumi Nagahama, Takahiro Okabe, Moulid Hidayat, Yuzo Kodama, Hideya Kitamura, Takashi Ogura, Norikazu Kitamura, Kazuho Ikeo, Shinichi Sasaki, Shigeru Tominaga, Kazuhisa Takahashi

## Abstract

Pirfenidone, an antifibrotic agent used for treatment of idiopathic pulmonary fibrosis (IPF), functions by inhibiting myofibroblast differentiation, which is involved in transforming growth factor (TGF)-β1-induced IPF pathogenesis. However, unlike normal lung fibroblasts, the relationship between pirfenidone responses of TGF-β1-induced human fibrotic lung fibroblasts and lung fibrosis is unknown. Here, we investigated the effect of pirfenidone on the functions of two new targets, collagen triple helix repeat containing protein 1 (CTHRC1) and four-and-a-half LIM domain protein 2 (FHL2), which included fibroblast activity, collagen gel contraction, and migration toward fibronectin. Compared to control lung fibroblasts, pirfenidone restored TGF-β1-stimulated fibroblast-mediated collagen gel contraction, migration, and CTHRC1 release in lung fibrotic fibroblasts. Furthermore, pirfenidone attenuated TGF-β1- and CTHRC1-induced fibroblast activity, bone morphogenic protein-4/Gremlin1 upregulation, and α-smooth muscle actin, fibronectin, and FHL2 downregulation, similar to that observed post-CTHRC1 inhibition. In contrast, FHL2 inhibition suppressed migration and fibronectin expression but did not downregulate CTHRC1. Overall, pirfenidone suppressed fibrotic fibroblast-mediated fibrotic processes via inverse regulation of CTHRC1-induced lung fibroblast activity. Thus, CTHRC1 can be used for predicting pirfenidone response and developing new therapeutic target for lung fibrosis.

**Summary statement:** Pirfenidone suppressed TGF-β1-mediated fibrotic processes in fibrotic lung fibroblasts by attenuating CTHRC1 expression, suggesting that CTHRC1 may be a novel therapeutic target for treating patients with lung fibrosis.

## Introduction

Accumulation of activated lung myofibroblasts and excessive deposition of extracellular matrix (ECM) produced by these cells result in lung tissue contraction, as has been observed in fibrotic lung tissues (Upagupta et al., 2018). This can disrupt lung function, and therefore, inhibition of fibrotic processes may alter the progression of lung fibrosis-related diseases. Diverse mediators, including Krebs von den Lungen (KL)-6 and surfactant protein (SP)-D released from damaged epithelial cells, and inflammatory cytokines (Aono et al., 2012;Xu et al., 2013) and pro-fibrotic growth factors (e.g., transforming growth factor [TGF]-β1 and platelet-derived growth factor [PDGF]) secreted by infiltrated inflammatory cells under conditions of airway inflammation induce fibrosis via autocrine mechanisms involving local lung fibroblast activation in the interstitial alveolar septa (Yoshida et al., 1992; Wynn, 2008).

TGF-β1, a key mediator of normal tissue repair (Blobe et al., 2000), strongly stimulates mesenchymal cells to produce large amounts of ECM, including fibronectin and collagen, resulting in the development of fibrosis (Yoshida et al., 1992). In addition, TGF-β1 stimulates fibroblast chemotaxis toward fibronectin (Sugiura et al., 2006;Togo et al., 2008) and augments fibroblast-mediated contraction of ECM by stimulating contractile stress fibers (α-smooth muscle actin [α-SMA]) (Kobayashi et al., 2006), generating lung fibroblasts that can be used as *in vitro* model of lung fibrosis. Fibronectin released from lung fibroblasts is a known autocrine or paracrine mediator of lung fibroblast-dependent chemotaxis and collagen gel contraction (Kamio et al., 2007;Togo et al., 2008).

Previous reports have shown that corticosteroid treatment does not improve prognosis in patients with idiopathic pulmonary fibrosis (IPF) (Nagai et al., 1999), suggesting that antifibrotic agents may be more useful than anti-inflammatory agents for the treatment of IPF. Pirfenidone (5-methyl-1-phenyl-2-(1H)-pyridone) is a potent antifibrotic agent that can inhibit the progression of fibrosis in patients with IPF. Pirfenidone attenuates the expression of procollagen, TGF-β1, and PDGF at the transcriptional level and ameliorates bleomycin-induced lung fibrosis in a rodent model (Iyer et al., 1999a; Iyer et al., 1999b;Gurujeyalakshmi et al., 1999). However, the precise mechanisms via which pirfenidone suppresses lung fibrosis are still unclear.

In this study, we evaluated the effects pirfenidone on TGF-β1-mediated contraction of ECM and migration toward fibronectin of lung fibroblasts isolated from patients with lung fibrosis and compared them with those of normal lung fibroblasts for understanding the mechanisms mediating lung fibroblast-dependent antifibrotic effects of pirfenidone. In addition, we focused on the two molecular targets of pirfenidone, namely, collagen triple helix repeat containing protein 1 (CTHRC1) and four-and-a-half LIM domain protein 2 (FHL2); the levels of these two proteins were previously demonstrated to be attenuated in the bleomycin-induced lung fibrosis model and CTHRC1 secretion was inhibited in the TGF-β1–induced normal primary human lung fibroblasts (Bauer er al.,2015). Our results provided important insights into pirfenidone-mediated antifibrotic processes. Furthermore, we established novel evidence of clinical markers for predicting responses to pirfenidone, which will assist in selecting therapy based on *in vitro* functional measurements of lung fibrotic fibroblast and clinicopathological information formally evaluated using multidisciplinary diagnosis (MDD).

## Results

### Clinical and demographic characteristics

The clinical and demographic characteristics of the patients are shown in Table 1. The two groups were similar in terms of age, smoking status, and sex. However, they differed significantly in lung function; as expected, patients with lung fibrosis had a lower percent forced vital capacity (% FVC). Histological examination revealed that of 12 patients with lung fibrosis not receiving medication, six had nonspecific interstitial pneumonia (NSIP), and six had usual interstitial pneumonia (UIP). Clinical diagnoses revealed three patients with IPF, five patients with NSIP, and four patients with chronic hypersensitivity pneumonitis (CHP).

**Table 1.** Clinical and demographic characteristics of the patients

| | No. of control patients | Mean ± SD | No. of IP patients | Mean ± SD | *P* value |
| --- | --- | --- | --- | --- | --- |
| Age, years | 12 | 64.5 ± 8.1 | 12 | 59.1 ± 14.7 | 0.28 |
| Sex, no. male/female | 9/3 |  | 8/4 |  | 1.00 |
| Smoking history (yes/no) | 8/4 |  | 8/4 |  | 1.00 |
| Pack-years | 8 | 712.5 ± 374.4 | 8 | 537.5 ± 498.2 | 0.44 |
| % FVC | 12 | 99.2 ± 4.1 | 12 | 85.2 ± 4.8 | 0.04 |
| KL-6 | 12 | None | 12 | 1422.3 ± 595.0 | |
| SP-D | 12 | None | 12 | 200.7 ± 77.8 | |
| Clinical diagnosis | 12 | None | 3 IPF, 5 NSIP, 4 CHP* |  |  |
| Histological pattern | 12 | None | 6 UIP, 6 NSIP* |  |  |

Abbreviations: CHP, chronic hypersensitivity pneumonitis; FVC, forced vital capacity; KL: Krebs von den Lungen; SP: surfactant protein; IP: interstitial pneumonia; IPF, idiopathic pulmonary fibrosis; NSIP, nonspecific interstitial pneumonia; UIP, usual interstitial pneumonia. \*, Diagnosed by multidisciplinary diagnosis (MDD).

### Effects of pirfenidone on TGF-β1-stimulated fibroblast activity

Pirfenidone inhibited collagen gel contraction of human fetal lung fibroblast-1 (HFL-1) in a concentration-dependent manner, but did not affect chemotaxis when added alone to HFL-1 cells (Fig. 1A, C).

**Fig. 1.**
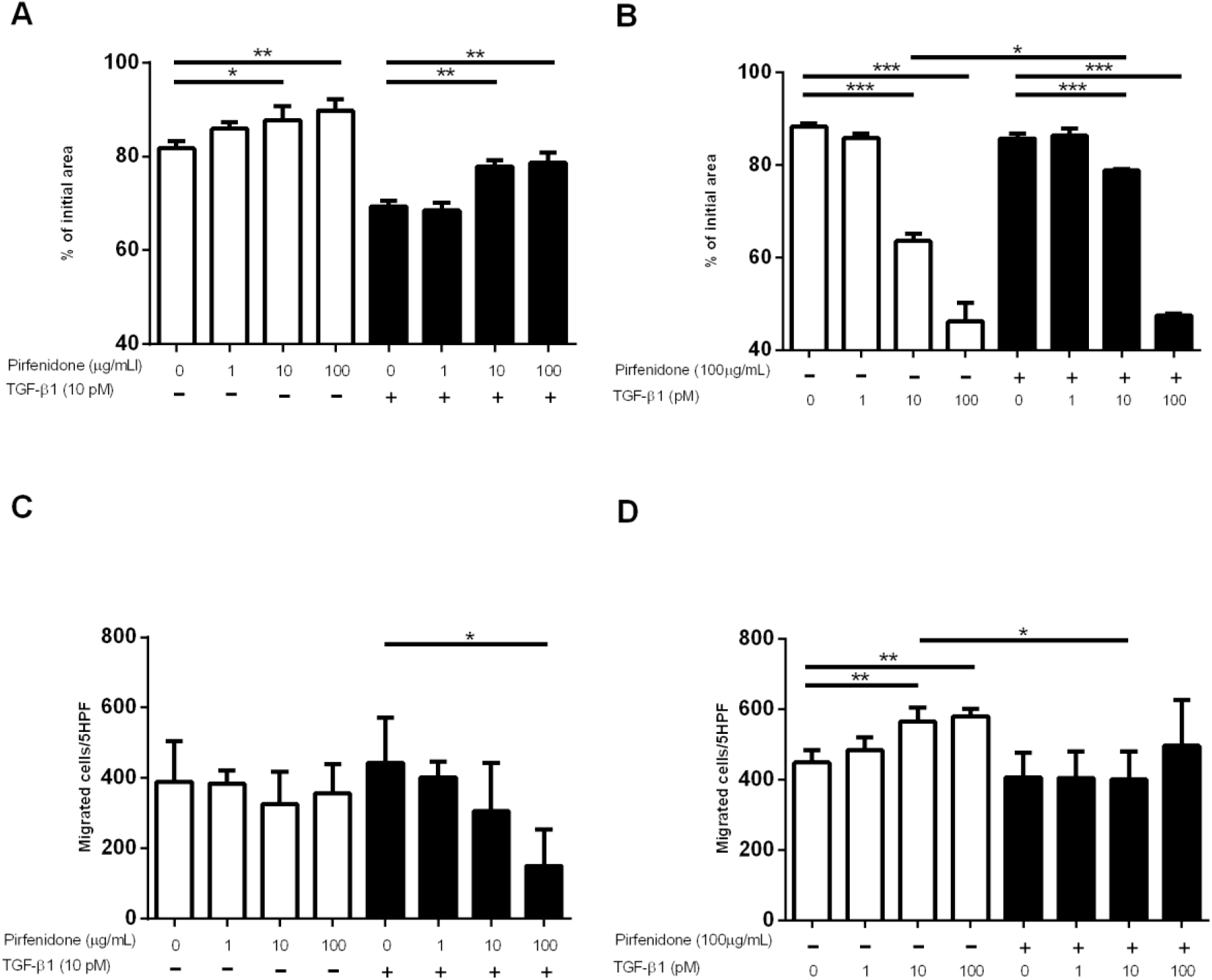
Effects of pirfenidone on TGF-β1-stimulated collagen gel contraction and chemotaxis in HFL-1 cells. HFL-1 cells were cultured and cast into three-dimensional collagen gels that were maintained in suspension, with gel size measured daily. HFL-1 cells also were grown in a monolayer culture and were trypsinized, and their chemotactic activity towards fibronectin (20 µg/mL) was assessed. (A) Collagen gel contraction on various concentrations of pirfenidone in the presence or absence of 10 pM TGF-β1. (B) Collagen gel contraction following treatment with various concentrations of TGF-β1 in the presence or absence of 100 μg/mL pirfenidone. (C) Number of migrated fibroblasts following treatment with various concentrations of pirfenidone in the presence or absence of 10 pM TGF-β1. (D) Number of migrated fibroblasts following treatment with various concentrations of TGF-β1 in the presence or absence of 100 μg/mL pirfenidone. Collagen gel contraction vertical axis: gel size measured after 2 days of contraction expressed as a percentage of the initial value. Chemotaxis vertical axis: number of migrated cells per five high-power fields (5 HPF). Horizontal axis: conditions. Values represent means ± SEMs of at least three separate experiments. Student’s unpaired t test. \**P* < 0.05, \*\**P* < 0.01, \*\*\**P* < 0.001.

Next, we investigated whether pirfenidone altered the TGF-β1-induced increase in collagen gel contraction and chemotaxis towards fibronectin in HFL-1 cells. Pirfenidone treatment reduced TGF-β1-induced collagen gel contraction and chemotaxis in HFL-1 cells in a concentration-dependent manner (*P* < 0.05 for 100 μg/mL pirfenidone ± 10 pM TGF-β1 versus control; Fig. 1A, C). Pirfenidone abolished HFL-1-mediated gel contraction and chemotaxis in the presence of less than 10 pM TGF-β1 (*P* < 0.05). However, in the presence of 100 pM TGF-β1, pirfenidone did not inhibit HFL-1-mediated gel contraction and chemotaxis (Fig. 1B, D).

Although the maximum plasma concentration (*C*_max_) of pirfenidone, which is typically used for treating patients with IPF at dosage up to 1800 mg/day, is 15.7 μg/mL after administration of 801 mg pirfenidone (Rubino et al., 2009), higher concentrations may be used; indeed, pirfenidone is widely used at concentrations above 100 μg/mL *in vitro* in the laboratory setting (Conte et al., 2014). Therefore, we used 100 μg/mL pirfenidone, with or without 10 pM TGF-β1, in our subsequent experiments on adult human primary lung fibroblasts.

Notably, gel contraction and chemotaxis were attenuated in cells treated with 100 μg/mL pirfenidone alone or in combination with 10 pM TGF-β1 (Fig. 2A, C). This inhibitory effect was higher in fibroblasts from fibrotic lungs than in control fibroblasts (gel contraction: *P*<0.001 and 0.05, chemotaxis: *P* < 0.05 and *P* =0.16, fibrotic vs. normal lung; Fig. 2B, D). However, there were no differences in the effects of fibrotic lung fibroblasts from patients with UIP compared to those from patients with NSIP. At the end of the incubation, cell numbers or viability in the gels in pirfenidone- and/or TGF-β1-treated groups were not different from those in the control group, as assessed by 3-(4,5-dimethylthiazol-2-yl)-2,5-diphenyltetrazolium bromide (MTT) assays (data not shown).

**Fig. 2.**
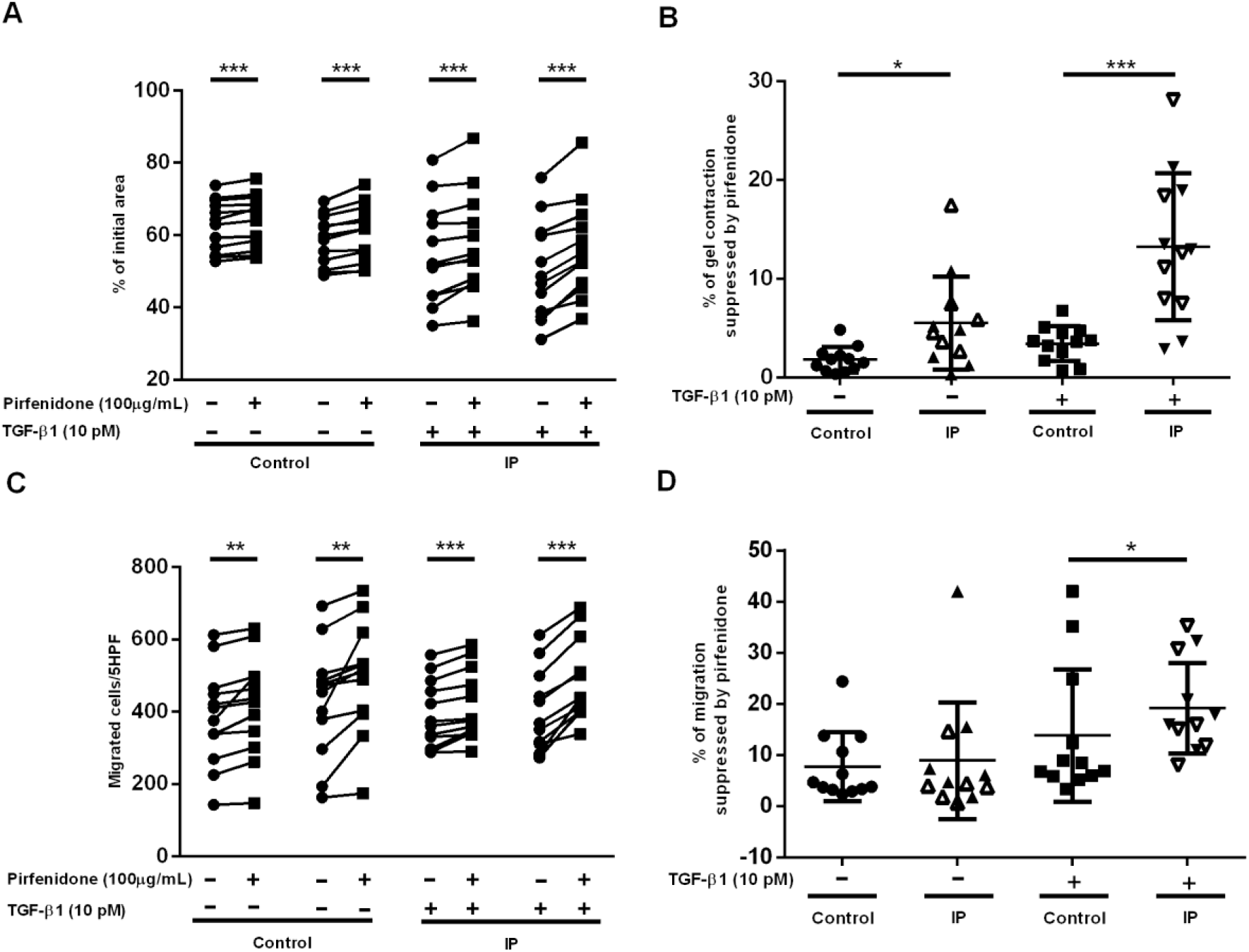
Effects of pirfenidone on collagen gel contraction and chemotaxis in lung fibroblasts from controls and patients with interstitial pneumonia (IP). Fibroblasts from controls and patients with IP were cultured in the presence or absence of pirfenidone (100 μg/mL) and 10 pM TGF-β1, and collagen gel contraction and chemotaxis were assayed. (A) Collagen gel contraction. Vertical axis: gel size measured after 2 days of contraction expressed as a percentage of the initial gel area. Horizontal axis: conditions. (B) Pirfenidone-induced suppression of collagen gel contraction in lung fibroblasts isolated from controls and patients with IP. Vertical axis: percentage of gel contracted size following pirfenidone treatment ([difference of gel size / initial gel size] × 100%). Horizontal axis: conditions. (C) Fibroblast chemotaxis. Vertical axis: number of migrated fibroblasts per 5 HPF. Horizontal axis: conditions. (D) Pirfenidone-dependent suppression of chemotaxis in lung fibroblasts isolated from controls and patients with IP. Vertical axis: percentage of migrated cells following pirfenidone treatment ([difference of migrated cells number / initial migrated cells number] × 100%). Horizontal axis: conditions. Each patient evaluated was expressed as an individual symbol, representing the means of at least two experiments each conducted in triplicate. (A, C) Lines connect the values for individual patients in the presence or absence of pirfenidone. \**P* < 0.05, \*\**P* < 0.01, \*\*\**P* < 0.001. (B, D) Filled symbols represent UIP, and open symbols represent NSIP. Student’s paired t test and unpaired t test. \**P* < 0.05, \*\**P* < 0.01, \*\*\**P* < 0.001.

### High sensitivity of TGF-β1-induced fibrotic mediators in fibrotic lung fibroblasts

CTHRC1 is a marker of activated stromal cells (Duarte et al., 2014). As pirfenidone suppresses both TGF-β1-induced CTHRC1 and FHL2 (Bauer et al., 2015), we assessed TGF-β1-induced changes in cytoplasmic CTHRC1 and FHL2 expression levels in fibrotic fibroblasts using immunoblotting. The relative increases in CTHRC1 and FHL2 upon TGF-β1 stimulation were higher in fibrotic fibroblasts than in control fibroblasts; however, there were no differences in fibrotic lung fibroblasts between patients with UIP and those with NSIP (CTHRC1: control, *P* = 0.109 versus lung fibrosis, *P* = 0.003; FHL2: control, *P* = 0.360 versus lung fibrosis, *P* = 0.006; Fig. 3A, B). Since CTHRC1 is a secreted protein and a previous report showed that CTHRC1 was present in plasma of individual patients (Duarte et al., 2014); therefore, we also measured release of CTHRC1 protein from human lung fibroblasts using enzyme-linked immunosorbent assay (ELISA). The relative increases in CTHRC1 releasing from fibroblasts upon TGF-β1 stimulation were higher from fibrotic fibroblasts than from control fibroblasts. The fold increase in CTHRC1 in the presence of TGF-β1 stimulation (1.347 ± 0.246 in control fibroblasts versus 3.610 ± 0.662 in fibrotic fibroblasts; *P* = 0.004) was higher in fibroblasts from fibrotic lungs than in control fibroblasts. The suppressive effects of pirfenidone on CTHRC1 release from TGF-β1-induced fibroblasts was higher in fibrotic lung fibroblasts than in control fibroblasts (control: *P* = 0.111; IP: *P* < 0.001); in contrast, the suppressive effects of pirfenidone alone were similar with respect to release of CTHRC1 (control: *P* = 0.004; IP: *P* = 0.001). The fold reduction in CTHRC1 in the presence of pirfenidone following TGF-β1 stimulation (0.139 ± 0.069 in control fibroblasts versus 0.400 ± 0.064 in fibrotic fibroblasts; *P* = 0.018) was higher in fibroblasts from fibrotic lungs than in control fibroblasts (All folds calculated by difference value after treatment / initial value) (Fig. 3C).

**Fig. 3.**
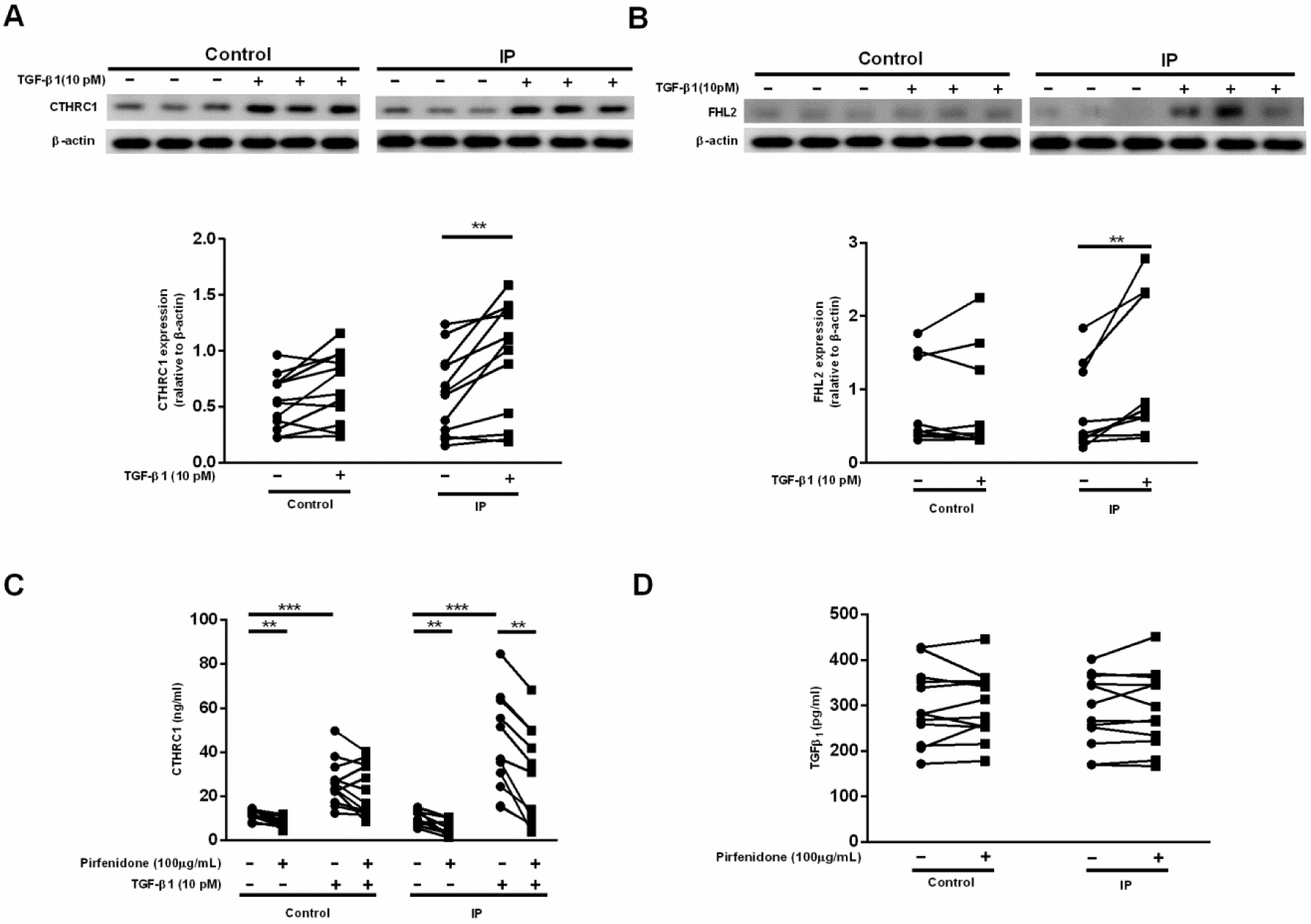
High sensitivity of TGF-β1-induced fibrotic mediators in fibrotic lung fibroblasts. Subconfluent fibroblasts from 12 controls and 12 patients with IP were cultured in serum-free (SF)-DMEM for 24 h and then incubated in the presence or absence of TGF-β1 (10 pM) and/or pirfenidone (100 ng/mL) for 48 h. Proteins from monolayer cultured fibroblasts were extracted and subjected to western blot analysis, and media were harvested from monolayer cultures and evaluated for CTHRC1 and TGF-β1 by immunoassay. Expression of (A) CTHRC1 (30 kDa) and (B) FHL2 (30 kDa) from fibroblasts isolated from controls and patients with IP in the presence or absence of TGF-β1 (10 pM). Vertical axis: expression of proteins normalized to expression of β-actin. Immunoassay of (C) CTHRC1 and (D) TGF-β1. Vertical axis: mediator production expressed as an amount. Symbols represent the mean values for individual patients, as assessed in two separate experiments. Horizontal axis: conditions. Student’s paired t test. \**P* < 0.05, \*\**P* < 0.01.

Fibroblasts are known to release mediators, including TGF-β1 (Sugiura et al., 2006; Togo et al., 2008) and prostaglandin E_2_ (PGE_2_) (Kohyama et al., 2001; Zhu et al., 2001), which can modulate chemotaxis and collagen gel contraction in an autocrine or paracrine manner. To determine whether these mediators directly contribute to the suppression of collagen gel contraction and chemotaxis induced by pirfenidone, the release of these mediators into the monolayer culture medium was evaluated using ELISA. Notably, pirfenidone did not affect TGF-β1 levels in culture medium of control and fibrotic fibroblasts (Fig. 3D). Furthermore, the ability of pirfenidone to modulate PGE_2_ release and inducible cyclooxygenase 2 (COX2) expression was further assessed in culture medium from HFL-1 cells. Pirfenidone did not stimulate PGE_2_ release or COX2 expression with or without TGF-β1 in fibroblasts (Fig. S1A, B). These results indicated that the pirfenidone-regulated molecules CTHRC1 and FHL2 may play a dominant role in pirfenidone-mediated fibroblast activity rather than exert direct effects on TGF-β1-mediated autocrine/paracrine regulation in fibroblasts.

### Effects of pirfenidone on TGF-β1-mediated fibrotic regulators in lung fibroblasts

Next, we assessed whether pirfenidone altered targets related to TGF-β1-mediated fibrotic processes in HFL-1 cells. Treatment with pirfenidone significantly reduced TGF-β1-augmented expression of CTHRC1, FHL2, α-SMA, fibronectin, and Gremlin1 (*P*<0.05; Fig. 4A–E, G) and reversed TGF-β1-dependent suppression of bone morphogenic protein-4 (BMP4; *P*<0.05; Fig. 4A, F).

**Fig. 4.**
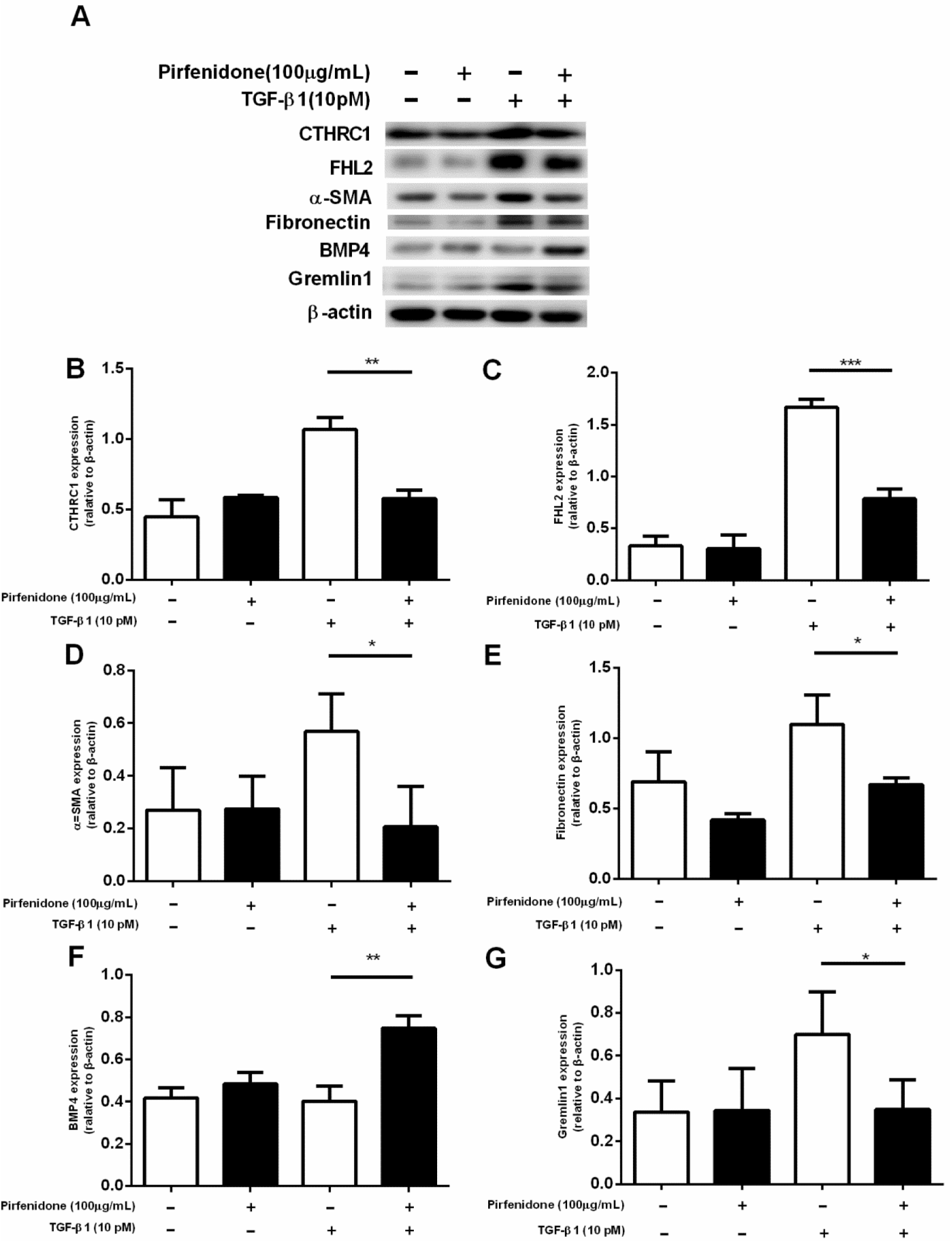
Effects of pirfenidone on TGF-β1-mediated fibrotic regulators in lung fibroblasts. Subconfluent HFL-1 cells were cultured in SF-DMEM for 24 h and then incubated in the presence or absence of TGF-β1 (10 pM) and pirfenidone (100 ng/mL) for 48 h. Total protein was extracted, and western blotting was performed with antibodies against the indicated proteins. (A) Western blot analysis of the effects of pirfenidone on targets related to TGF-β1-mediated fibrotic processes, including CTHRC1 (30 kDa), FHL2 (30 kDa), α-SMA (42 kDa), fibronectin (250 kDa), BMP4 (47 kDa), Gremlin1 (25 kDa), and β-actin (42 kDa). The vertical axis shows the relative intensities of (B) CTHRC1, (C) FHL2, (D) α-SMA, (E) fibronectin, (F) BMP4, and (G) Gremlin1 versus β-actin; the horizontal axis shows the conditions. Values represent means ± SEMs of at least three independent experiments. Student’s unpaired t test. \**P* < 0.05, \*\**P* < 0.01, \*\*\**P* < 0.001.

### Effects of pirfenidone on CTHRC1-mediated regulation in lung fibroblasts

We investigated the effects of recombinant human (rh) CTHRC1 on HFL1-mediated collagen gel contraction and chemotaxis. rhCTHRC1 (10–1000 ng/mL) stimulated gel contraction and chemotaxis toward fibronectin in a concentration-dependent manner (*P* < 0.05) compared to the control (Fig. 5A–C), accompanied by concentration-dependent upregulation of FHL2, α-SMA, fibronectin, and Gremlin1 and downregulation of BMP4 (Fig. 5A). According to a previous report on detection of CTHRC1 plasma levels, we used 100 ng/mL rhCTHRC1 with or without 10 pM TGF-β1 in our studies on HFL-1cells (Duarte et al., 2014). Pirfenidone significantly attenuated rhCTHRC1-induced gel contraction and chemotaxis (*P* < 0.05; Fig. 5E, F); it also suppressed rhCTHRC1-augmented FHL2, α-SMA, fibronectin, and Gremlin1 expression (*P*<0.05; Fig. 5D–I, K) and reversed rhCTHRC1-dependent suppression of BMP4 (*P*<0.05; Fig. 5D, J).

**Fig. 5.**
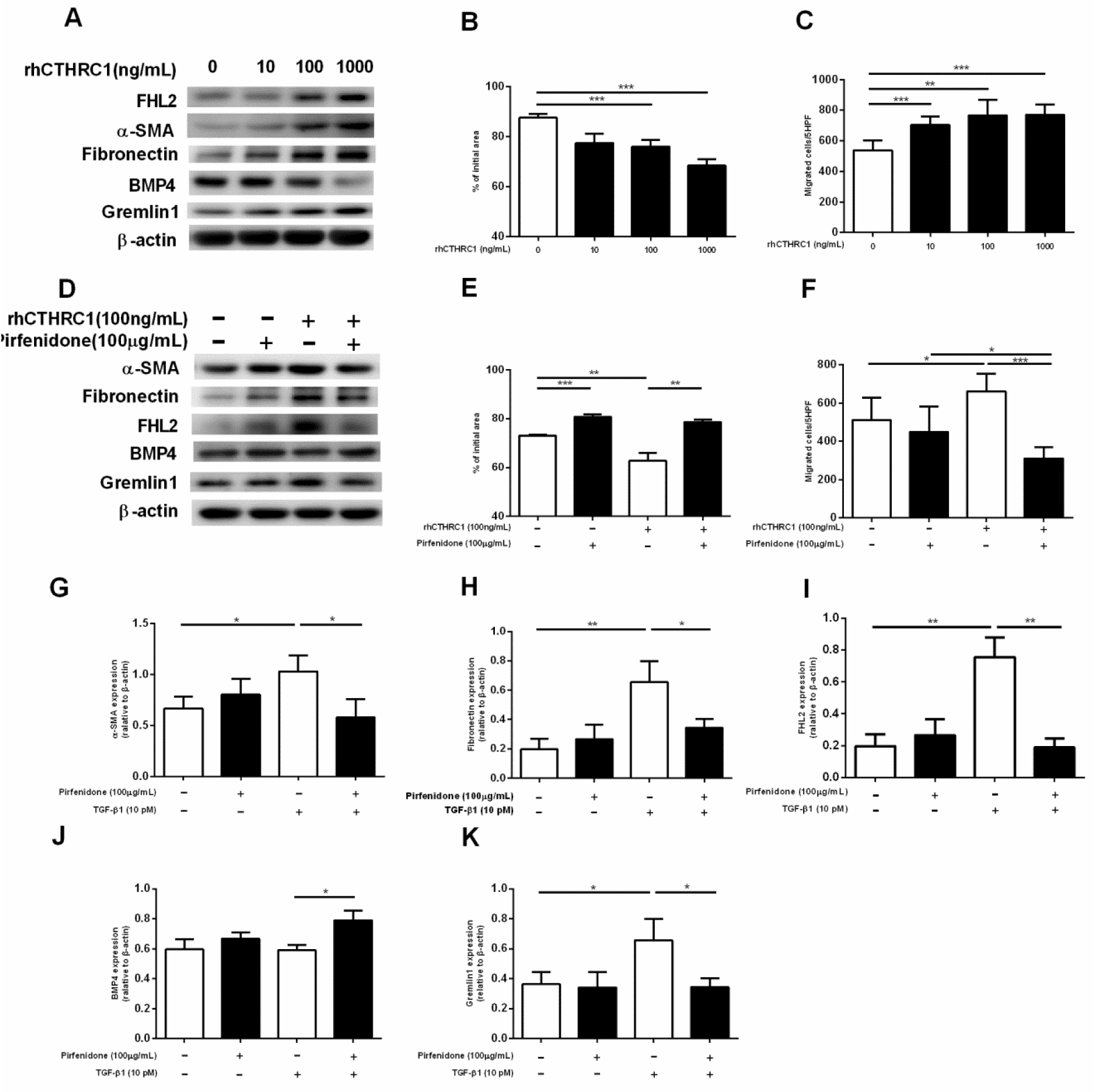
Effects of pirfenidone on CTHRC1-mediated regulation in lung fibroblasts. Subconfluent HFL-1 cells were cultured in SF-DMEM for 24 h and then incubated in the presence or absence of different concentrations of rhCTHRC1 (see Methods). (A) Western blot analysis of the effects of different concentrations of rhCTHRC1 on targets related to fibrotic processes, i.e., FHL2 (30 kDa), α-SMA (42 kDa), fibronectin (250 kDa), BMP4 (47 kDa), Gremlin1 (25 kDa), and β-actin (42 kDa). Effects of different concentrations of rhCTHRC1 on HFL-1 cell-mediated collagen gel contraction (B) and chemotaxis (C). Subconfluent HFL-1 cells were cultured in SF-DMEM for 24 h and then incubated in the presence or absence of rhCTHRC1 (100 ng/mL) and pirfenidone (100 ng/mL) for 48 h. (D) Western blot analysis of the effects of pirfenidone on rhCTHRC1-mediated targets related to fibrotic processes, i.e., FHL2 (30 kDa), α-SMA (42 kDa), fibronectin (250 kDa), BMP4 (47 kDa), Gremlin1 (25 kDa), and β-actin (42 kDa). Effects of pirfenidone on rhCTHRC1-mediated collagen gel contraction (E) and chemotaxis (F). Collagen gel contraction, vertical axis: gel size measured after 2 days of contraction expressed as a percentage of the initial value. Chemotaxis, vertical axis: number of migrated cells per 5 HPF. Horizontal axis: conditions. Effects of pirfenidone on the expression levels of rhCTHRC1-mediated targets assayed using western blot analysis. The vertical axis shows the relative intensities of (G) α-SMA, (H) fibronectin, (I) FHL2, (J) BMP4, (K) Gremlin1 versus β-actin; the horizontal axis shows the conditions. Values represent means ± SEMs of at least three independent experiments. Student’s unpaired t test. \**P* < 0.05, \*\**P* < 0.01, \*\*\**P* < 0.001.

### Inhibition of CTHRC1- and FHL2-mediated regulation of HFL-1

To investigate the functional roles of the pirfenidone-mediated targets CTHRC1 and FHL2 in lung fibroblasts, we knocked down these targets in HFL-1 cells. Silencing of *CTHRC1* led to a complete reversal of targets related to TGF-β1-mediated fibrotic processes, i.e., reduction in FHL2, α-SMA, fibronectin, and Gremlin1 expression and increase in BMP4 expression (Fig. 6A). Furthermore, *CTHRC1* knockdown attenuated gel contraction and chemotaxis (Fig. 6B, C). However, silencing of *FHL2* did not affect CTHRC1, α-SMA, BMP4, and Gremlin1 expression and gel contraction, but reduced fibronectin expression and chemotaxis (Fig. 6D, E).

**Fig. 6.**
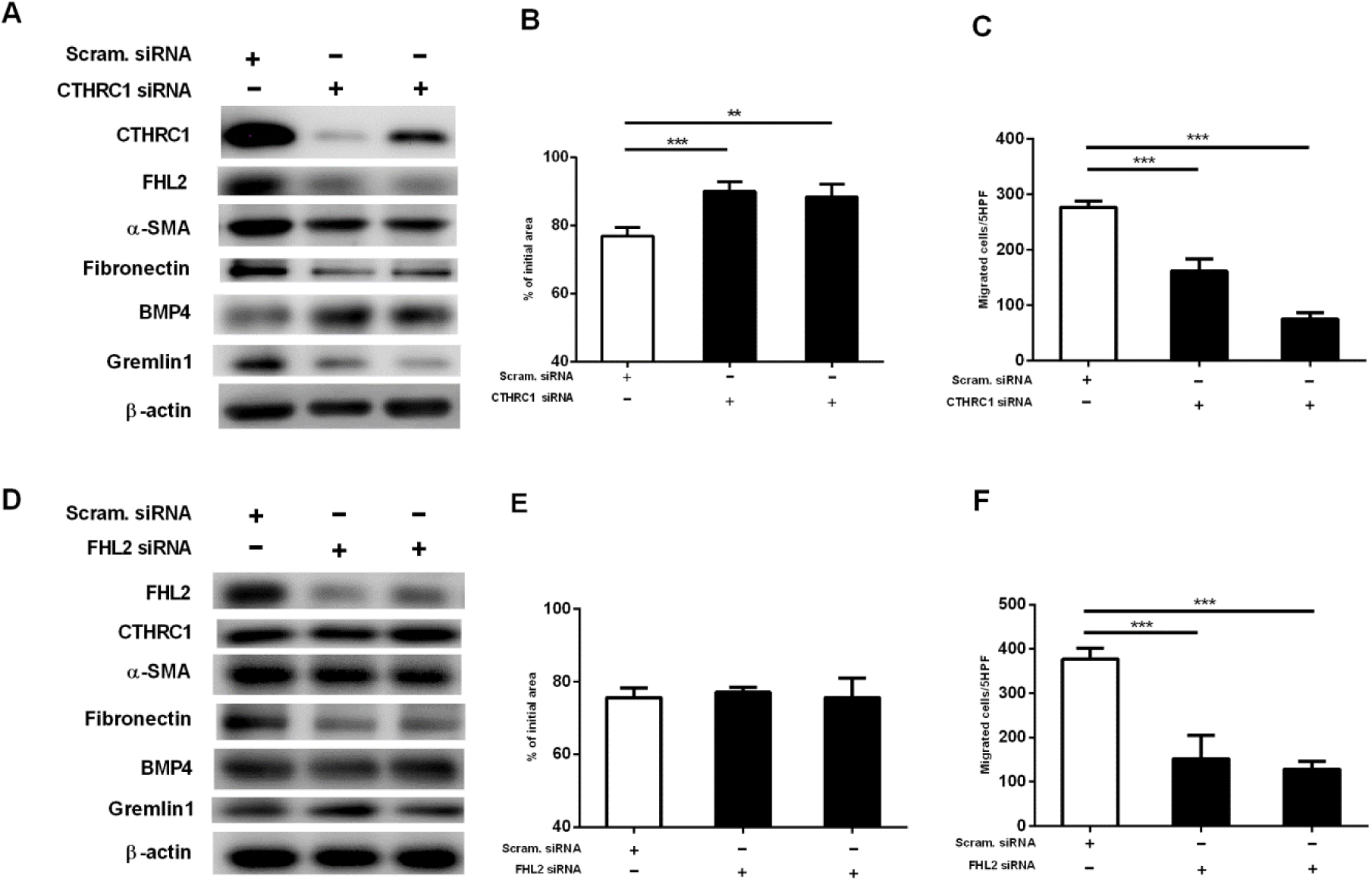
Effects of *CTHRC1* and *FHL2* knockdown in HFL-1 cells. *CTHRC1*-and *FHL2*-knocked down HFL-1 cells were examined for collagen gel contraction and chemotaxis. (A) Western blot analysis of the effects of *CTHRC1* silencing on targets related to fibrotic processes, i.e., CTHRC1 (30 kDa), FHL2 (30 kDa), α-SMA (42 kDa), fibronectin (250 kDa), BMP4 (47 kDa), Gremlin1 (25 kDa), and β-actin (42 kDa). (B) Collagen gel contraction and (C) chemotaxis after silencing of *CTHRC1*. (D) Western blot analysis of the effects of *FHL2* silencing on targets related to fibrotic processes, i.e., FHL2 (30 kDa), CTHRC1 (30 kDa), α-SMA (42 kDa), fibronectin (250 kDa), BMP4 (47 kDa), Gremlin1 (25 kDa), and β-actin (42 kDa). (E) Collagen gel contraction and (F) chemotaxis after silencing of *FHL2*. Collagen gel contraction, vertical axis: gel size measured after 2 days of contraction expressed as a percentage of the initial value. Chemotaxis, vertical axis: number of migrated cells per 5 HPF. Horizontal axis: conditions. Values represent means ± SEMs of at least three separate experiments. Student’s unpaired t test. \**P* < 0.05, \*\**P* < 0.01, \*\*\**P* < 0.001.

### Effects of pirfenidone on *in vitro* TGF-β1-stimulated fibroblast activity and biomarkers of lung fibrosis

Considering our observation that pirfenidone in fibrotic lung fibroblasts enhanced their sensitivity to TGF-β1-induced fibrosis compared to normal fibroblasts, we investigated whether our *in vitro* data was clinically significant. Toward this objective, we measured serum levels of the lung fibrosis biomarkers KL-6 and SP-D in serum of patients with lung fibrosis at the time of primary lung fibroblast sample collection. The ability of pirfenidone to abrogate the TGF-β1-induced increase in collagen gel contraction correlated positively with SP-D levels (r^2^ = 0.388, *P* = 0.031, Fig. 7B) but not with KL-6 levels (Fig. 7A). In contrast, the inhibitory effects of pirfenidone on TGF-β1-induced migration correlated negatively with KL-6 levels (r^2^ = 0.380, *P* = 0.033, Fig. 7C) but not with SP-D levels (Fig. 7D). No other clinical or histopathological and spirometric parameters, such as sustainer and rapid decliner categories according to forced vital capacity (FVC) reduction rates, were related to *in vitro* fibroblast response to pirfenidone treatment or to *in vitro* fibroblast activity.

**Fig. 7.**
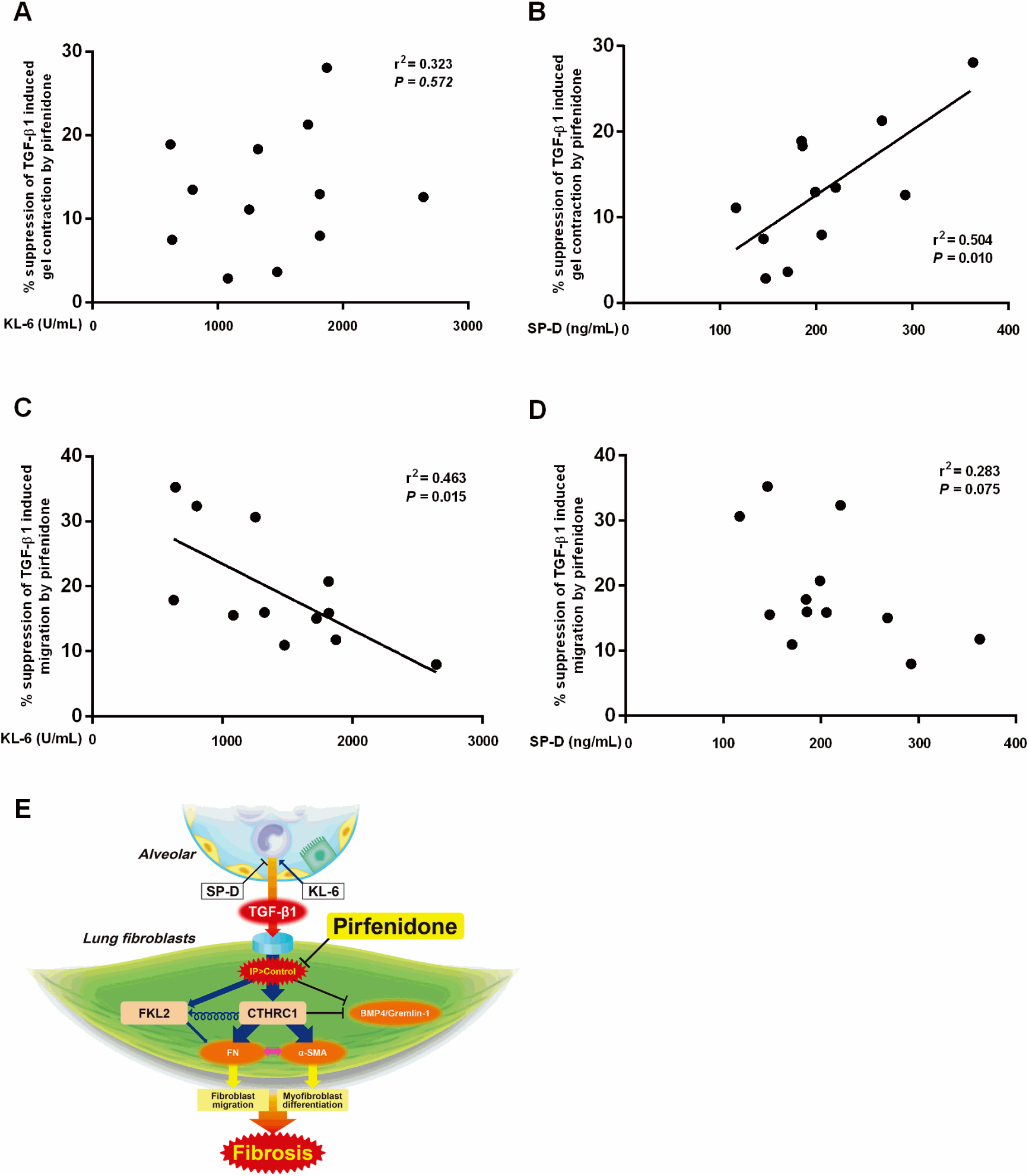
Relationship between pirfenidone responses to TGF-β1-stimulated fibroblast bioactivity in vitro and biomarkers of lung fibrosis. Comparison of the relationships between suppression of TGF-β1-induced gel contraction by pirfenidone and serum (A) KL-6 and (B) SP-D levels and between suppression of TGF-β1-induced migration and (C) KL-6 and (D) SP-D levels. Symbols represent individual patients. Linear regression. *P* < 0.05 indicates a positive relationship between pirfenidone response to TGF-β1-stimulated fibroblast bioactivity in vitro and biomarkers of lung fibrosis. (E) Through airway cell/fibroblast interaction, lung fibroblasts are continually exposed to TGF-β1, which is regulated by mediators released by airway cells under inflammatory conditions (Aono et al., 2012) (Xu et al., 2013), resulting in distinct phenotype of fibrotic fibroblasts. This high sensitivity to TGF-β1 along with upregulation of CTHRC1 and FHL2 leads to the development of fibrosis in fibrotic fibroblasts. Treatment with pirfenidone can effectively block this process.

## DISCUSSION

In this study, we observed that changes in the fibrotic lung fibroblast phenotype resulted in increased response to the pirfenidone-suppressed bioactivity of lung fibroblasts, leading to inhibition of collagen gel contraction and migration, following TGF-β1-mediated activation. Previous reports have shown that pirfenidone reduces proliferation, migration, fibroblast-embedded collagen gel contraction, ECM production, and TGF-β1-mediated differentiation into myofibroblasts by attenuating the effect of TGF-β1 and its downstream targets, including phosphorylation of Smad3, connective tissue growth factor, p38, and Akt in human fibroblasts (Conte et al., 2014; Hall et al., 2018; Lin et al., 2009;Saito et al., 2012;Shi et al., 2011;;Sun et al., 2018), which support our current results. Pirfenidone prevented changes in the fibrotic fibroblast phenotype, increasing proliferation and migration and enhancing levels of phospho-Smad3, phospho-signal transducer and activator of transcription 3, α-SMA, and collagen in the context of IPF (Epstein Shochet et al., 2018). We clarified the mechanisms via which CTHRC1 and FHL-2 (Bauer et al., 2015} were stimulated by TGF-β1 in lung fibrotic fibroblasts compared to in control fibroblasts. Our results showed that pirfenidone suppressed CTHRC1-induced migration toward to fibronectin, gel contraction, and α-SMA and fibronectin expression and increased the BMP4/Gremlin1 ratio. In additionr, pirfenidone also suppressed FHL-2-mediated fibroblast migration.

CTHRC1 is expressed by activated stromal cells of diverse origin and co-expressed with α-SMA. Elevated CTHRC1 levels were detected in patients with inflammatory conditions, including rheumatoid arthritis, and CTHRC1 is considered a marker of tissue remodeling, inflammation, or wounding (Duarte et al., 2014; Shekhani et al., 2016). A previous report has shown that CTHRC1 plays a protective role in pulmonary fibrosis and tissue repair and may be clinically applied for treating fibrosis as it decreases collagen matrix deposition by inhibiting Smad2/3 activation (LeClair and Lindner, 2007). In addition, rhCTHRC1 inhibits TGF-β1-stimulated collagen type I synthesis and promotes skin repair in keloid fibroblasts (Li et al., 2011; Pyagay et al., 2005). The systemic loss of CTHRC1 increased TGF-β1-mediated excess matrix deposition and induced the development of bleomycin-induced lung fibrosis in mice (Binks et al., 2017). Furthermore, TGF-β1 and BMP-4, which belong to the TGF-β superfamily of ligands, stimulate CTHRC1 expression, and overexpression of CTHRC1 in smooth muscle cells, and embryonic fibroblasts treated with exogenous rhCTHRC1 increases migration (Pyagay et al., 2005; Shekhani et al., 2016). In this study, we demonstrated that fibrotic fibroblasts showed altered phenotypes of TGF-β1-stimulated CTHRC1 expression and pirfenidone-dependent suppression of CTHRC1 expression after TGF-β1 treatment. In addition, rhCTHRC1 stimulated lung fibroblast-mediated gel contraction and migration, and *CTHRC1* knockdown suppressed fibronectin expression, migration toward fibronectin, and gel contraction. Pirfenidone restored TGF-β1- and rhCTHRC1-dependent suppression of BMP4 expression, which was previously reported to reduce lung fibroblast proliferation and TGF-β1-induced synthesis of ECM components, including fibronectin (Jeffery et al., 2005; Pegorier et al., 2010). In the present study, pirfenidone also reduced the levels of the BMP4 antagonist Gremlin1, which was previously implicated in the development of lung fibrosis (Myllarniemi et al., 2008), when stimulated by TGF-β1 and rhCTHRC1. These results indicated that CTHRC1 acted as a costimulator of TGF-β1-mediated fibrotic processes in fibrotic fibroblasts rather than as a suppresser of fibrosis. Thus, CTHRC1 could be a dominant target for pirfenidone-mediated antifibrotic mechanisms in fibrotic lung fibroblasts.

FHL2 participates in tissue wound healing and is associated with fibrogenesis (Gullotti et al., 2011; Wixler et al., 2007). *Fhl2* ^−/-^ mice develop hepatic fibrogenesis (Dahan et al., 2017), and *Fhl2*-deficient embryonic fibroblasts show reduced collagen contraction and cell migration, resulting in impaired wound healing (Wixler et al., 2007). However, in contrast to CTHRC1, FHL2 has been reported to positively regulate the expression of collagen types I and III in an FHL2-induced BLM-treated lung fibrosis model; this process tended to involve acceleration of lung inflammation rather than direct FHL2-induced fibrotic mechanisms (Alnajar et al., 2013; Kirfel et al., 2008; Park et al., 2008). Furthermore, FHL2 induces α-SMA (Wixler et al., 2007) and is stimulated by TGF-β1 (Bauer et al., 2015; Muller et al., 2002}. In this study, we also demonstrated that FHL2 was further stimulated by TGF-β1 in fibrotic fibroblasts compared to control fibroblasts and that pirfenidone suppressed TGF-β1-stimulated FHL2 expression. However, *FHL2* knockdown suppressed only fibronectin expression and migration toward fibronectin, but did not affect gel contraction, other TGF-β1-mediated fibroblast regulators, or CTHRC1 expression. As *CTHRC1* knockdown attenuated FHL2 expression, FHL2 may play a role in lung fibroblast-mediated fibrosis following TGF-β1-induced upregulation of *CTHRC1* (Fig. 7E).

Clinically, patients with IPF who showed predicted vital capacity (VC) of more than 70% and lowest oxygen satuation in blood during 6 min walking tests (less than 90% at baseline) are most likely to benefit from pirfenidone therapy (Azuma et al., 2011). The expression levels of pirfenidone-targeted translational gene markers (*GREM1, CTHRC1*, and *FHL2*) showed significant negative correlation with the percentage diffusing capacity of carbon monoxide (%DLCO), and was associated with IPF disease severity (Bauer et al., 2015). However, little is known regarding the surrogate markers of pirfenidone response. In this study, we demonstrated that the magnitude of pirfenidone-dependent suppression of TGF-β1-induced gel contraction and migration was positively related to serum SP-D levels and negatively related to serum KL-6 levels, but was not related to any other clinical parameters, including histological pattern and lung function (%VC, %FVC, and %DLCO). These diverse phenotypic responses to pirfenidone in fibrotic lung fibroblasts and our *in vitro* results related to serum biomarkers suggested that the progressive state of lung fibrosis with increasing SP-D/KL-6 ratios may be associated with better clinical outcomes (Fig. 7E). This analysis described the relative usefulness of other clinical parameters at baseline when estimating the predictable surrogate marker of patients with lung fibrosis as candidates for pirfenidone therapy.

The results presented here provided evidence regarding the high sensitivity of fibrotic fibroblasts to pirfenidone, which mediated TGF-β1-induced fibrotic processes. However, our study has certain limitations. For example, we used limited number of patients’ fibroblasts lines and did not assess plasma levels of CTHRC1 as a clinical surrogate marker for predicting pirfenidone response. Instead, we performed functional experiments in lung fibroblasts and analyzed the responses of human lung fibrosis patient-derived lung fibroblast samples to pirfenidone for determining the utility of CTHRC1 as a potential therapeutic target and for predicting pirfenidone responses in lung fibroblast-mediated fibrosis regulation. Our observations provide preliminary evidence regarding identification of pirfenidone responses in personalized therapies and the application of CTHRC1 as a novel biomarker with translational value for predicting pirfenidone responses in patients with lung fibrosis.

## MATERIALS AND METHOD

### Materials

Cell culture medium (Dulbecco’s modified Eagle’s medium [DMEM]) was purchased from Wako (Osaka, Japan). Fetal calf serum was purchased from Sigma-Aldrich (St. Louis, MO, USA). TGF-β1 was obtained from R & D Systems (Minneapolis, MN, USA), and rhCTHRC1 was from Abcam (Cambridge, UK). Pirfenidone was provided by Shionogi & Co. Ltd. (Osaka, Japan) and was dissolved in 100% dimethylsulfoxide (DMSO). The amount of DMSO added did not affect the results of the bioassays (Hatzelmann and Schudt, 2001). Preliminary experiments with MTT demonstrated that the concentrations of pirfenidone and DMSO used in this study did not show any significant cytotoxicity in fibroblasts (data not shown).

### Cell culture

HFL-1 cells were obtained from the American Type Culture Collection (CCL-153; Manassas, VA, USA). Primary lung fibroblasts were obtained from 12 patients with lung fibrosis, as diagnosed by a multidisciplinary team using the gold standard approach (Chung and Lynch, 2016), and 12 patients without clinical airway symptom or lung functional abnormalities as a control group (Table 1). The Institutional Review Board of Juntendo University School of Medicine and Kanagawa Cardiovascular and Respiratory Center approved the procedures. All patients provided written, informed consent (approval no. 2012173). Human primary lung parenchymal fibroblasts from patients undergoing lung resection were cultured as described previously (Holz et al., 2004). Tissues were obtained from areas of macroscopically normal lung parenchyma, distal to any tumor masses, in the control fibroblast group. Cells were collected from areas of complete fibrosis (resembling a honey comb), avoiding large airways, vessels and pleural surface in the lung fibrosis fibroblast group. These cells were cultured as described above. Cells from the outgrowths of these cells, termed “P0,” were frozen for later use; these cells displayed typical fibroblast morphology and were confirmed to be positive for vimentin and negative for cytokeratin. For chemotaxis, three-dimensional collagen gel contraction, and ELISA analyses, primary lung fibroblasts were used at passages 4–6 after isolation to exclude the effects of differences in passage number and culturing conditions.

### Fibroblast chemotaxis

HFL-1 cell chemotaxis was assessed using the Boyden blindwell chamber technique (Neuroprobe, Inc., Gaithersburg, MD, USA) as previously described (Boyden, 1962). In experiments with TGF-β1 or rhCTHRC1, pirfenidone was added to the wells of the upper chamber, whereas human fibronectin (20 µg/mL) was placed in the bottom chamber as the chemoattractant. The two wells were separated by an 8-µM pore filter (Nucleopore, Pleasanton, CA, USA). The chambers were incubated at 37°C in a humid atmosphere containing 5% CO_2_ for 8 h, after which the cells on top of the filter were removed by scraping. The cells that had migrated through the filter were then fixed, stained with DiffQuick Sysmex (16920), and mounted on glass microscope slides. Migration was assessed by counting the number of cells in five high-power fields. Replica experiments were performed in triplicate, and replicates with separate cell cultures were performed on separate occasions. Wells with serum-free DMEM were used as negative controls.

### Collagen gel contraction assay

Type I collagen (rat tail tendon collagen) was extracted from rat tail tendons as previously described (Elsdale and Bard, 1972). The effects of pirfenidone on fibroblast-mediated gel contraction were determined in the presence or absence of TGF-β1 or rhCTHRC1 using a modification of the method developed by Bell et al. (Bell et al., 1979). The floating gels were cultured for up to 3 days, and the ability of the fibroblasts to contact the gels was determined by quantifying the area of the gels daily using an LAS4000 image analyzer (GE Healthcare Bio-Science AB, Uppsala, Sweden). Data are expressed as the gel area percentage compared to the original gel size.

### Measurement of CTHRC1, TGF-β1, and PGE_2_ levels

Cultures were maintained for 48 h to quantify CTHRC1, TGF-β1, and PGE_2_ levels. After 48 h, media were collected, frozen, and stored at −80°C until analysis. CTHRC1, TGF-β_1_, and PGE_2_ production by the cells was determined using human CTHRC1 (LifeSpan BioScienceds, Inc., Seattle, WA, USA), TGF-β_1_ (R&D Systems), and PGE_2_ immunoassays (Cayman Chemical, Ann Harbor, MI, USA), respectively, according to the manufacturers’ instructions.

### Western blot analysis

To standardize culture conditions, cells were passaged at a density of 5 × 10^4^ cells/mL, cultured for 48 h, and then collected for preparation of whole cell lysates. The medium was changed to DMEM without serum for 24 h, followed by treatment with TGF-β1 (10 pM) or rhCTHRC1 (100 ng/mL) in the presence or absence of pirfenidone (100 μg/mL) for 48 h or with various concentrations of rhCTHRC1 for 8 h. Primary antibodies against the following proteins were used for western blotting: CTHRC1 (1:5000 dilution; Proteintech, Rosemont, IL, USA; cat. no. 16534-1-AP), FHL2 (1:1000 dilution; Abcam, Cambridge, UK; cat. no. ab66399), α-SMA (1:1000 dilution; Sigma-Aldrich; cat. no. A2547), fibronectin (1:1000 dilution; Enzo Life Sciences, Inc., Farmingdale, NY, USA; cat. no. BML-FG6010-0100), BMP-4 (1:1000 dilution; Abcam; cat. no. ab39973), Gremlin1 (1:1000 dilution; Thermo Fisher Scientific, Waltham, MA, USA; cat. no. PA5-13123), β-actin (1:5000 dilution; Wako Pure Chemical Industries; cat. no. 013-24553), and COX2 (1:1000 dilution; Abcam; cat. no. ab169782). Bound antibodies were visualized using peroxidase-conjugated secondary antibodies and enhanced chemiluminescence with a LAS4000 image analyzer (GE Healthcare Bio-Science AB), and band intensity was analyzed Image Gauge software (LAS-400 Plus; Fujifilm).

### Small interfering RNA (siRNA)-mediated knockdown assays

Commercial siRNA targeting CTHRC1 (10620318; Life Technologies, Carlsbad, CA, USA) and FHL2 (1027416; Qiagen, Valencia, CA, USA) was transfected using RNAiMAX transfection reagent (13778-150; Life Technologies) diluted in Opti-MEM (31985062; Gibco/Life Technologies) according to the manufacturers’ instructions. HLF-1 cells were plated at 1 × 10^5^ cells/mL and incubated for 24 h, and were used for transfection when they reached 50–70% confluency. Predetermined concentrations of siRNA were used to achieve more than 70% knockdown. To suppress endogenous CTHRC1 and FHL2 in fibroblasts, the cells were transfected for 24 h with 50 nM CTHRC1 siRNA or 15 nM FHL2 siRNA. A scrambled siRNA probe was used as a control. After silencing CTHRC1 or FHL2 with siRNA, the cells were analyzed using western blotting, and collagen gel contraction assays and chemotaxis experiments were performed.

### Statistical analysis

Results are expressed as means±standard errors of the means (SEMs). We used Student’s t tests to determine differences between groups and linear regression to relationship. For experiments in which paired samples within a group were available, we used paired Student’s t tests. For these comparisons, each patient was considered an individual data point. Differences with *P* values less than 0.05 were considered significant. Data were analyzed using Prism 6 software (GraphPad Inc., San Diego, CA, USA).

## Acknowledgments

We thank X. Liu, F. Sakai, T. Takemura, T. Iwasawa, I. Taki, L. Ning, N. Okamoto, T. Ito, R. Mineki, F. Takahashi, and E. Kobayashi for technical assistance and advice. We thank T. Takeda and N. Kondo for providing us CAGE result to clarified mechanism from RIKEN Center for Life Science Technologies. We thank Tadayuki Takeda and Naoto Kondo for providing us CAGE result try to further clarify present mechanism from RIKEN Center for Life Science Technologies. We thank S. Kusano and A. Kusano for drawing figure 7E design from KUSANO DESIGN OFFICE (Tokyo, Japan). And we acknowledges the International Fellowship Program, Takeda Science Foundation.

## Abbreviation list

α-SMA: α-smooth muscle actin
BMP-4: bone morphogenetic protein-4
CHP: chronic hypersensitivity pneumonitis
COX2: cyclooxygenase 2
CTHRC1: collagen triple helix repeat containing protein 1
%DLCO: percent diffusing capacity of carbon monoxide
ECM: extracellular matrix
ELISA: enzyme-linked immuno sorbent assay
FHL2: four-and-a-half LIM domain protein 2
%FVC: percent forced vital capacity
HFL-1: human fetal lung fibroblast-1
IP: interstitial pneumonia
IPF: idiopathic pulmonary fibrosis
KL: Krebs von den Lungen
MDD: multidisciplinary diagnosis
NSIP: nonspecific interstitial pneumonia
PDGF: platelet-derived growth factor
PGE2: prostaglandin E2
Rh: recombinant human
SEM: standard errors of the means
SP: surfactant protein
TGF: transforming growth factor
UIP: usual interstitial pneumonia
VC: vital capacity

## Supplement figure

**Fig. S1.**
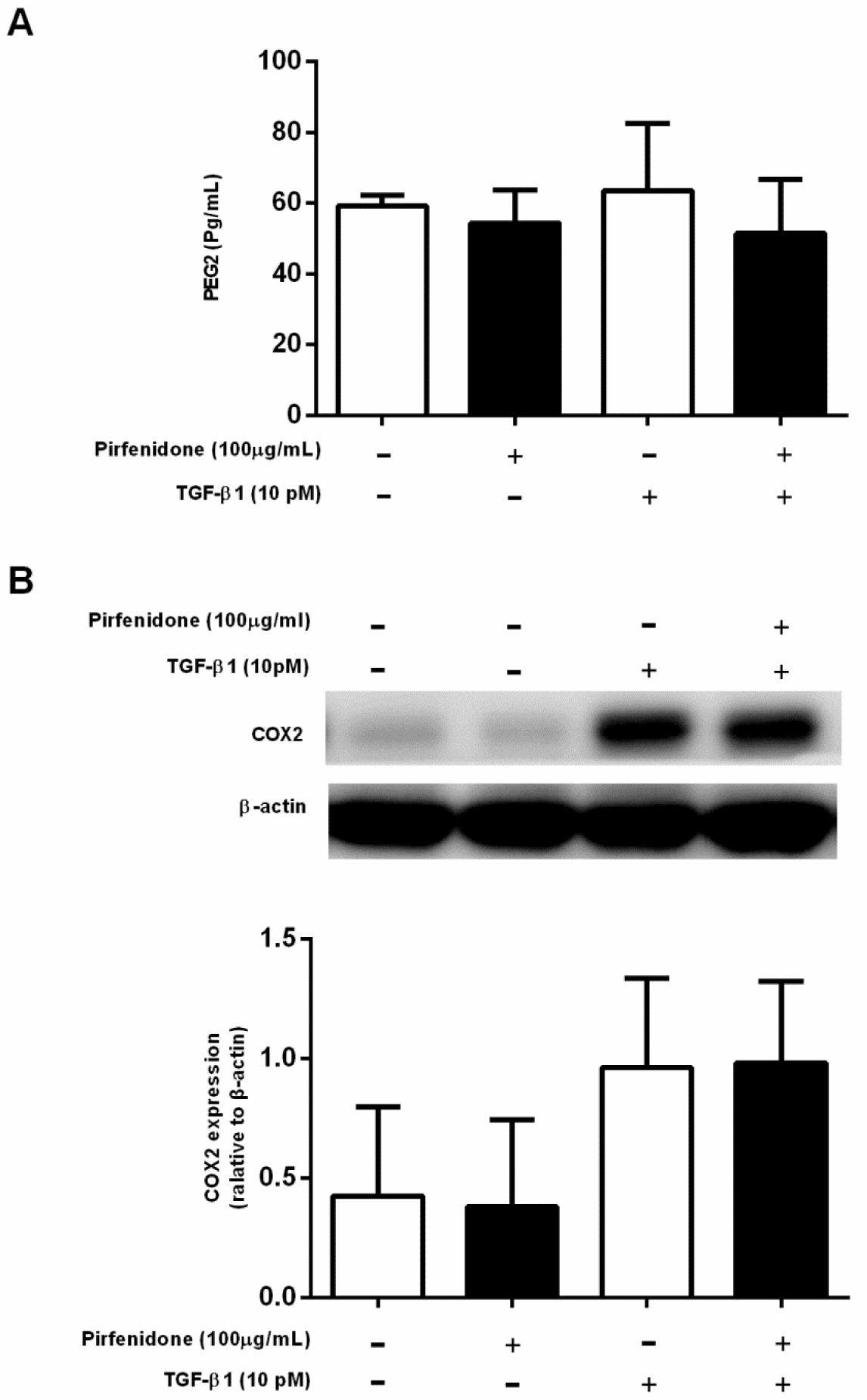
Effects of pirfenidone on TGF-β1-mediated regulators of the cyclooxygenase 2 (COX2)/prostaglandin E2 (PGE2) pathway in fibroblasts. Subconfluent HFL-1 cells were cultured in SF-DMEM for 24 h and then incubated in the presence or absence of TGF-β1 (10 pM) and pirfenidone (100 ng/mL) for 48 h. Proteins were extracted and subjected to western blot analysis of COX2. Media were harvested from monolayer culture and evaluated for PGE2 by immunoassay. (A) Release of PGE2 in HFL-1 monolayer cultures and (B) western blot analysis of COX2 (69 kDa). Bar figure vertical axis: protein expression relative to β-actin. Horizontal axis: conditions. Values represent means ± SEMs of at least three independent experiments.

